# Cell projection plots: a novel visualization of bone marrow aspirate cytology

**DOI:** 10.1101/2022.12.06.519348

**Authors:** Taher Dehkharghanian, Youqing Mu, Catherine Ross, Monalisa Sur, H.R. Tizhoosh, Clinton JV Campbell

**Affiliations:** McMaster University, Hamilton, Canada; Artificial Intelligence and Informatics, Mayo Clinic, Rochester, MN, USA; Juravinski Hospital and Cancer Centre, Hamilton, Canada

## Abstract

Deep models for cell detection have demonstrated utility in bone marrow cytology, showing impressive results in terms of accuracy and computational efficiency. However, these models have yet to be implemented in the clinical diagnostic workflow. Additionally, the metrics used to evaluate cell detection models are not necessarily aligned with clinical goals and targets. In order to address these issues, we introduce cell projection plots (CPPs), which are novel, automatically generated visual summaries of bone marrow aspirate specimens. CPPs provide a compact summary of bone marrow aspirate cytology, and encompass relevant biological patterns such as neutrophil maturation. To gauge clinical relevance, CPPs were shown to three hematopathologists, who decided whether shown diagnostic synopses matched with generated CPPs. Pathologists were able to match CPPs to the correct synopsis with 85% accuracy. Our finding suggests CPPs can compactly represent clinically relevant information from bone marrow aspirate specimens, and may be used to efficiently summarize bone marrow cytology to pathologists. CPP could be a step toward human-centered implementation of artificial intelligence (AI) in hematopathology, and a basis for a diagnostic support tool for digital pathology workflows.

## 1 Introduction

Artificial intelligence (AI), particularly deep networks, has shown promise in digital pathology [1]. AI models assist with information extraction from digital whole slide images (WSIs) of pathology specimens, supporting and automating various aspects of pathology diagnostic workflows [2]. However, AI models have not yet been widely implemented in the diagnostic hematopathology workflows [3]. In cytopathology, a number of AI models have been trained on bone marrow aspirate (BMA) images for cell detection and classification, including one from our group [4, 5, 6, 7, 8, 9]. These models often return a nucleated differential count (NDC) as their output, where many types of individual bone marrow cells are counted and classified based on subtle morphological features [10]. While important for a number of clinical decision points in hematology, the NDC has limited value in some hematological diagnoses such as myelodysplastic neoplasm (MDS), as it ignores the morphological complexities of BMA [11]. Additionally, such models do not relieve pathologists of the laborious task of viewing a large WSI to analyze thousands of cells. Therefore, in many cases, cell counts alone do not provide a summary of the complex information present needed to make a primary hematology diagnosis from BMA.

In digital pathology, AI models are often evaluated by measures such as accuracy, F1-score, and mean average precision (mAP); these are neither easy to interpret nor necessarily aligned with clinical problems and expectations in the medical field. Clinically meaningful evaluation of AI remains one of the challenges of its implementation in pathology [12]. Therefore, impressive performance in terms of computer science metrics does necessarily translate into a model’s clinical utility. In pathology practice, the ultimate goal of cytopathology examination is not to find the most accurate bounding boxes in detection, but rather to glean biologically relevant information from a specimen to make a diagnosis that predicts biology, or patient outcomes. The necessity of clinical evaluation of medical AI has been well-emphasized [13].

In the context of these limitations in practical utility and clinically-relevant evaluations, we present an efficient method to visualize cytological information extracted from BMA by a deep network to hematopathologists. We combined deep feature extraction and dimensionality reduction to create a compact visual summary of BMA slides, called a *cell projection plot* (CPP). Deep features are numerical vectors generated by using deep neural networks, which have been extensively used for image representation in both cytology and histopathology [14, 15, 16, 17, 18]. Dimensionality reduction has an established usage in visual analytics[19]. Therefore, a combination of deep feature extraction and dimensionality reduction is an intuitive approach to represent BMAs to pathologists in a user-friendly and interpretable way. In our previous work, we have described a system for detection and classification of BMA cells [5].

Here, we developed CPPs by a) re-purposing our published cell detection and classification model to serve as a feature extractor; b) applying dimensionality reduction to visualize detected cells on a 2D plane; c) evaluating the descriptive quality of CPPs by pathologists. Figure 1 shows an overview of the approach implemented in this study. Our results suggest CPP can serve as an automatically generated visual summary of BMA cytology, capturing biologically and diagnostically relevant information. Therefore, CPPs provide additional insight into bone marrow cytology beyond NDC, and may eventually contribute to the development of AI-based compact representations for primary diagnosis in digital hematopathology workflows.

**Figure 1:**
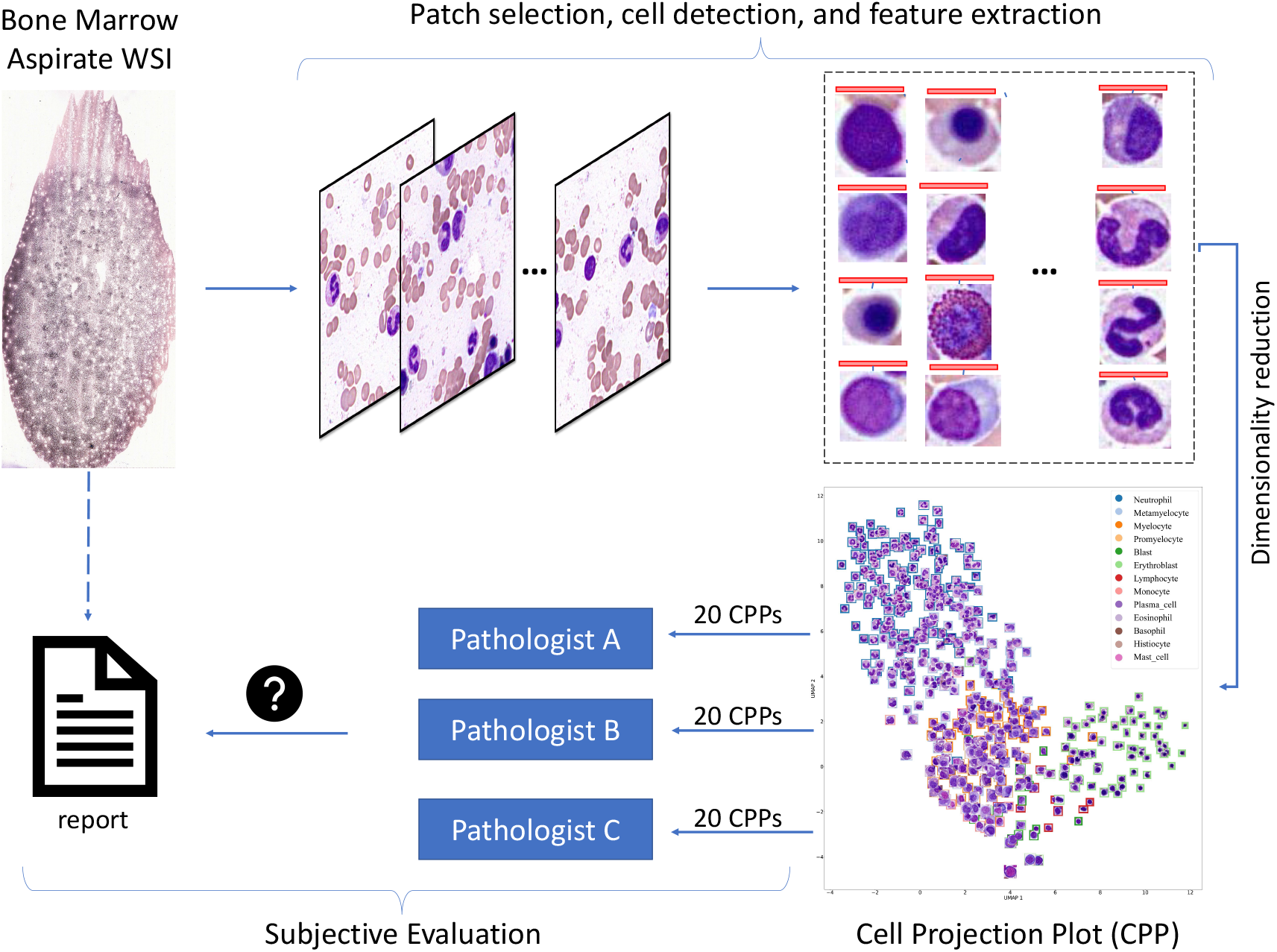
Overview of our approach. First, bone marrow aspirate WSI tiles were analyzed by a deep network (YOLO) for cell detection and feature extraction. Then, deep feature vectors were embedded in a two-dimensional space, and cells were visualized at their corresponding coordinates, to create CPPs. Three pathologists evaluated 60 CPPs (20 each), in order to assess if CPPs could provide diagnostically relevant information.

## 2 Materials and Methods

The College of American Pathologists (CAP) has published guidelines for validating digital imaging workflows for diagnostic purposes [20]. In this study, we adapted these guidelines for evaluating CPP as a means for workflow augmentation in hematopathology. Accordingly, we had three hematopathologists evaluate CPP samples of 20 patients each; 60 BMA samples in total.

### 2.1 Data acquisition, slides, patients

Sixty May–Grünwald–Giemsa stained bone marrow aspiration WSIs were used in this study. These WSIs were scanned at 40X and acquired in TIFF file format under Hamilton Research Ethics Board (HIREB) study protocol 7766-C. A custom software was developed to allow pathologists to view and evaluate WSIs and CPPs.

### 2.2 Cell detection and feature extraction

A pre-trained YOLO object detection model for bone marrow cell detection was used in this study [5]. YOLO detects and classifies bone marrow cells from bone marrow aspiration digital WSI. This model is composed of three parts: backbone, neck, and head. Feature extraction takes place in the backbone. The backbone’s first output is used to produce deep feature vectors to represent each detected cell. Since detected cells are of different sizes, average pooling was applied to the corresponding regions to create a linear vector of size 256. This process is schematically depicted in Figure 2.

**Figure 2:**
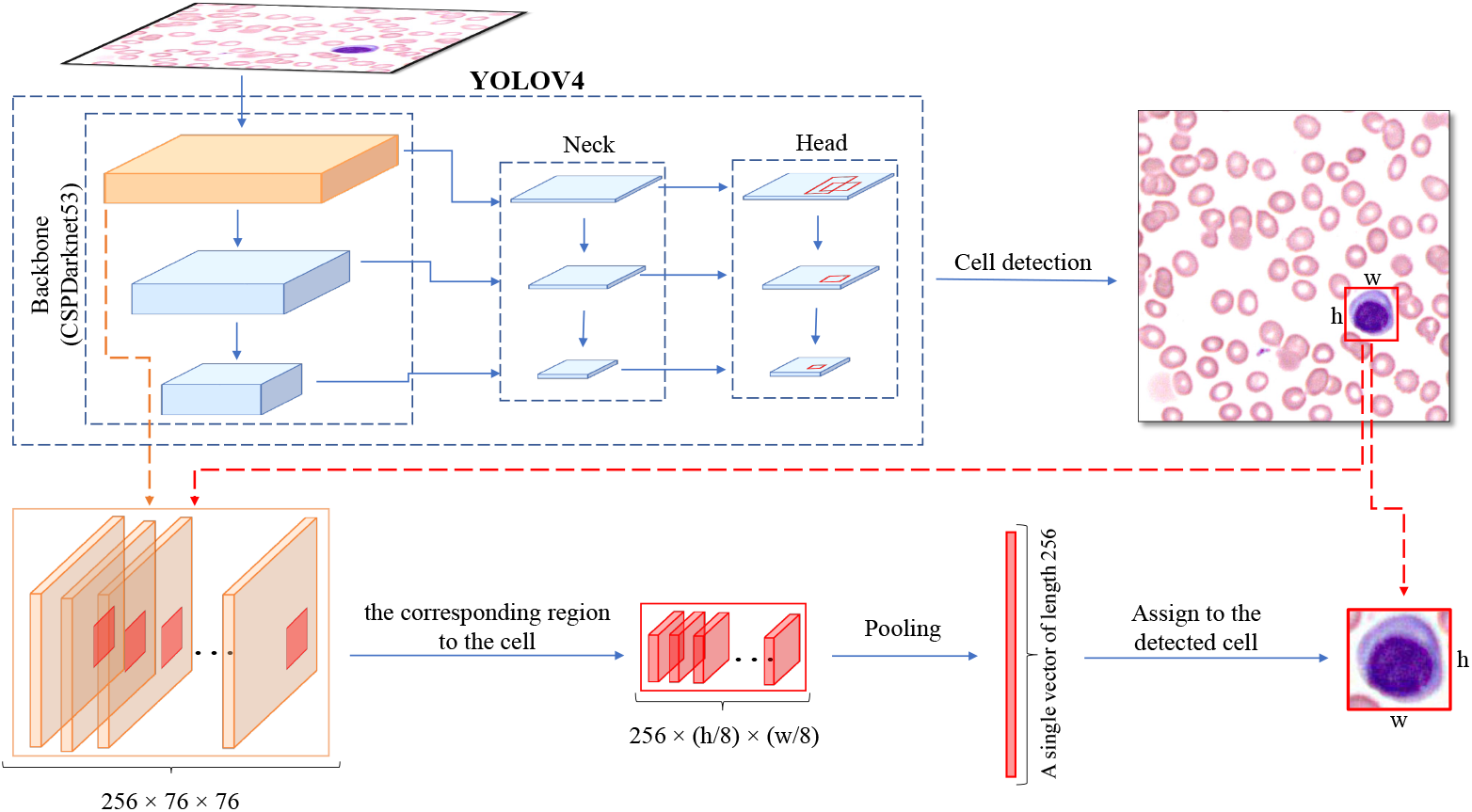
Overview of the feature extraction process. Tissue tiles were analyzed by a YOLO model to detect cells and extract deep features. A deep feature vector was produced for each detected cell from the corresponding region on the feature map, i.e., the first output of the model’s backbone. Average pooling was applied on the corresponding region to construct a deep feature vector of size 256 for each detected cell. These deep feature vectors represent cells.

### 2.3 Dimensionality Reduction

Uniform Manifold Approximation and Projection (UMAP) is a dimensionality reduction technique that preserves more of the global structure of the data compared to another popular dimensionality reduction algorithm, namely t-SNE [21]. UMAP was used to embed cell-specific deep feature vectors into two dimensions for visualization.

### 2.4 Cell Projection Plots

Although the object detection model detects 19 classes, only the following cells were included in this study, “blasts”, “promyelocytes”, “myelocytes”, “metamyelocytes”, “neutrophils”, “erythroblasts”, “lymphocytes”, “monocytes”, “plasma cells”, “eosinophils”, “basophils”, and “megakaryocytes”. Tissue tiles were sampled from digital bone marrow slides; the YOLO model was applied to tissue tiles to detect and classify cells, using the same pipeline described in our previous work [5]. 500 cells were sampled out of the detected cells in proportion to each cell type’s prevalence. Megakaryocytes were gathered by a modified method. Since megakaryocytes are not as abundant as other cell types and are often found in thick regions not useful for the NDC, more tissue tiles are needed to find one megakaryocyte. Therefore, all tissue tiles in a WSI were used to detect an adequate number of megakaryocytes. The same YOLO model was used for this purpose.

#### 2.4.1 Particles

We designed an algorithm to find particles from WSI thumbnail images. We started by converting each thumbnail to a single-channel grayscale picture, which is essential for the proper thresholding technique. We used Gaussian blurring to decrease the noise in this grayscale picture. After that, we employed thresholding to convert all pixels to binary: black and white, emphasizing the objects of interest, i.e., particles, for the contour-detection approach. In this binary image, we also used mathematical morphology operation *erosion* to eliminate artifacts. Finally, we used OpenCV’s [22] findContours function to determine the estimated contours of the particle. With particle contours in the thumbnail, we extracted particles from high-power images accordingly. Particles were shown to the pathologists so that they would have an estimate of the specimen’s cellularity.

### 2.5 Cell projection plot evaluation design

The experiment was designed to evaluate the usability of CPPs alone. In order to prevent confirmation bias, the process was slightly gamified. 60 CPPs were shown to three pathologists; each pathologist reviewed 20 CPPs. For each CPP, either the real synopsis reported or a randomly sampled report was shown to the pathologists. To check CPP’s descriptive quality, Pathologists were asked to decide whether a displayed synopsis report matched the generated CPP.

## 3 Results

### 3.1 Cell Projection Plots

The goal of dimensionality reduction is to shrink the high-dimensional feature space so that it can be visualized, while preserving the semantic relationship between individual embeddings. In order to assess the quality of the YOLO embeddings (representing cells) used for CPPs, we assessed a well-described biological process in hematopoiesis: granulocytic (neutrophil) maturation. To this end, we used UMAP [21] in cases of BMA WSI labeled as normal, acute myeloid leukemia (AML), myelodysplastic neoplasm (MDS) and chronic myeloid leukemia. These example cases represent different expected biological states in terms of granulocytic maturation. A typical example of normal maturation is shown in Figure 3a. This plot indicates a clear, continuous, and orderly granulocytic maturation pattern, with discernible neutrophil precursors subsets in expected biological proportion (blasts, promyelocytes, myelocytes, metamyelocytes, and neutrophils). The plot in Figure 4b is from an AML WSI, where could see that blasts dominate the plot, indicating maturation arrest. Figures 3c and 4d are generated from an MDS and a CML WSI. In these cases, there is continuous granulocytic maturation, with varying proportions of blasts (MDS) and intermediate myeloid precursors (CML) as expected in these disease states. This suggested that such visualizations could be useful for assessing semantically relevant patterns in BMA cytology, and that the YOLO embeddings were of good quality. Importantly, it indicated that visualizing deep cell embeddings by dimensionality reduction is biologically and clinically relevant.

**Figure 3:**
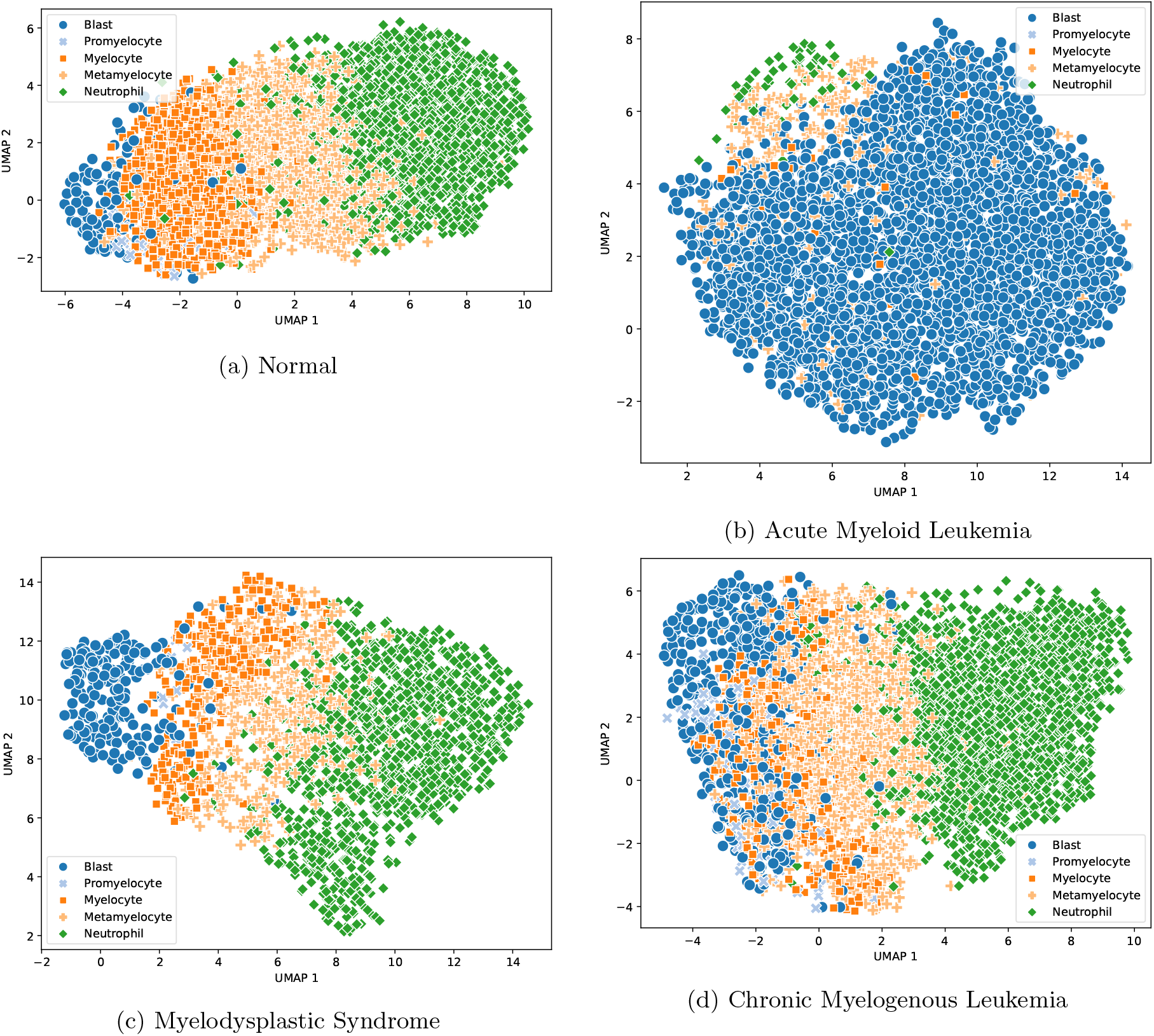
Four UMAP plots showing the maturation of neutrophils. Subfigure “a” shows orderly neutrophil maturation in a normal bone marrow aspirate WSI. The distribution of neutrophils and their precursors is in accordance with the expected biological maturation pattern, which suggests that the embeddings can be used to present semantically and clinically relevant patterns. Subfigure “b” is from an acute myeloid leukemia aspirate WSI where the blasts dominate the plot. Subfigure “c” is from a myelodysplastic neoplasm aspirate WSI. Subfigure “d” is from a chronic myelogenous leukemia aspirate WSI, where there are abundant intermediate myeloid precursors. Of note, UMAP *x* and *y* coordinate values are arbitrary.

**Figure 4:**
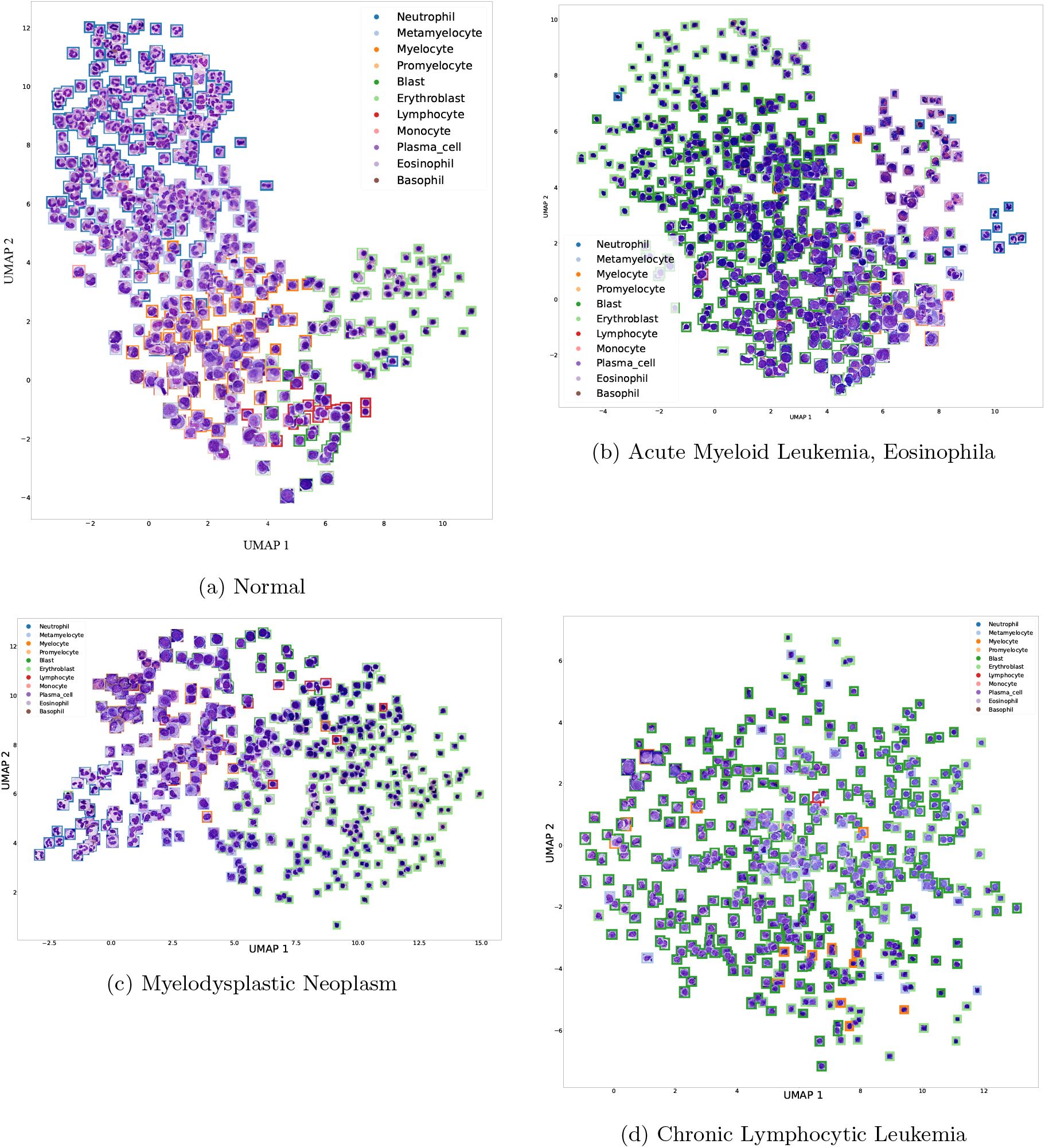
Four sample Cell Projection Plots (CPP). CPPs were evaluated by pathologists. Subfigure “a” is from a normal BMA indicating an orderly distribution of cells. Subfigure “b” shows an AML case with proportionally increased number of blasts and eosinophils. Subfigure “c” is from an MDS case (dysplastic erythropoiesis and granulopoiesis can be seen). Subfigure “d” shows a CLL case where the object detection model misclassified lymphocytes as blasts.

Next, we extended this concept to the entire cell population in a BMA, excluding megakaryocytes. CPPs are generated by adding individual cell images to the dimensionality reduction plots, thereby forming a compact representation of a BMA for visual assessment. Figure 4a shows a typical CPP of a normal BMA, as well as a variety of typical hematopoietic diseases. In the normal CPP, for example, it can be seen that granulopoiesis and erythropoiesis are present in acceptable relative proportions, with granulopoiesis showing continuous maturation. Granulopoiesis is a distinct population moving from the bottom left to the top left with maturation, while erythropoiesis is a distinct population to the middle right of the CPP.

Figure 4b shows a CPP of acute myeloid leukemia and eosinophilia BMA. The proportional increase in the number of blasts and eosinophils is evident. A CPP from myelodysplastic neoplasm shows an expansion of erythropoiesis and dysplastic myelopoiesis, as would be expected (Figure 4c), and a CPP for a chronic lymphocytic leukemia patient shows an expansion of abnormal lymphocytes (Figure 4d). Although the model misclassified large lymphocytes as blasts in this case, this could be recognized and intercepted by hematopathologists. Therefore, these findings suggested that pathologists can quickly review these CPPs as a compact representation of BMA cytology, and also, as an interpretable implementation of AI in BMA cytology.

Megakaryocytes were shown to hematopathologists in a CPP that did not include other cell types, due to their larger size and unique biological significance. Figure 5a shows a megakaryocyte CPP from a normal BMA, and Figure 5b is from an MDS patient, where hyposegmented megakaryocytes can be seen.

**Figure 5:**
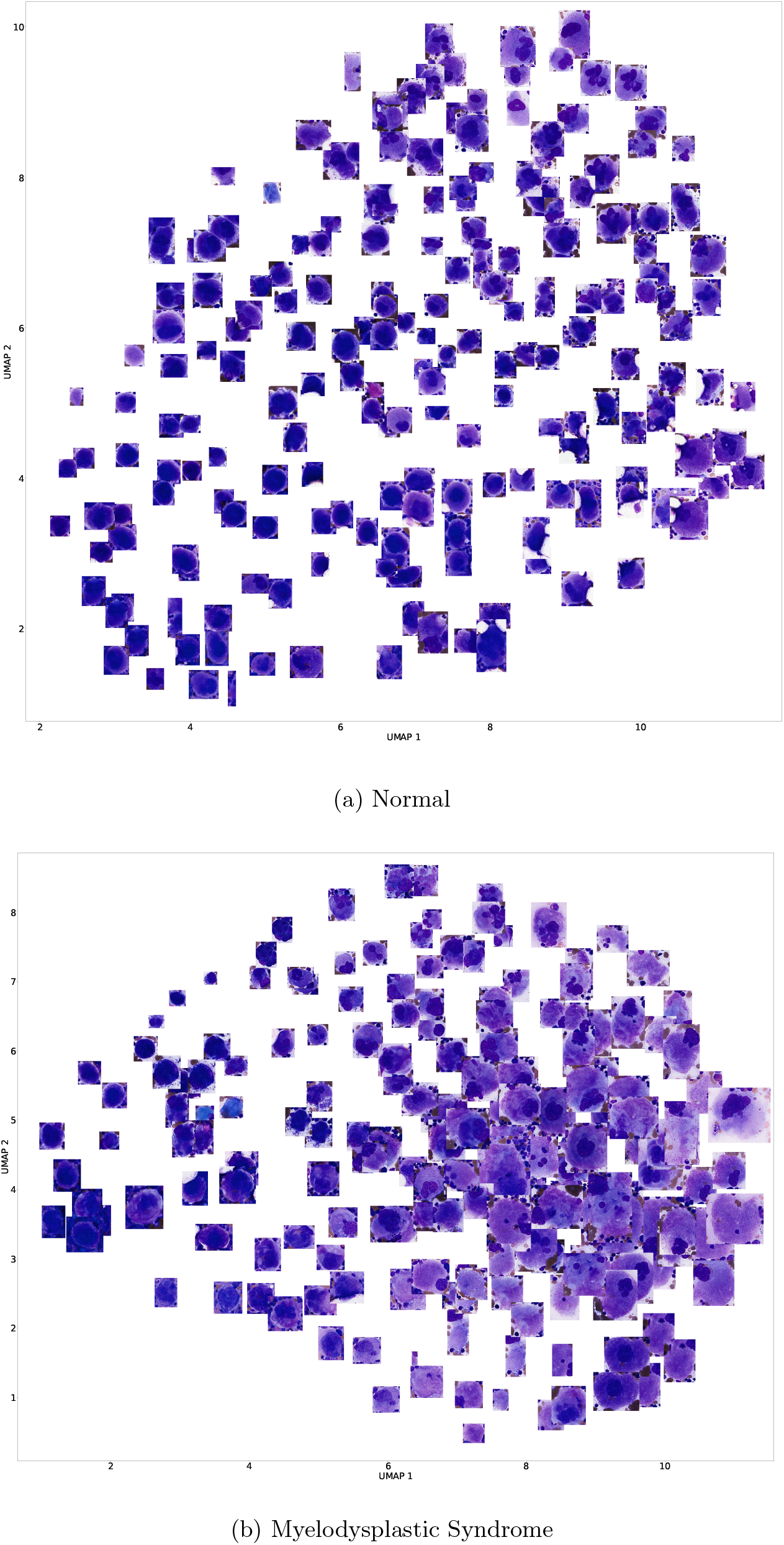
Two sample Cell Projection Plot (CPP) for megakaryocytes. CPPs were evaluated by pathologists. Subfigure “a” is from a normal BMA. Subfigure “b” shows an MDS case and hyposegmented megakaryocytes are can be seen.

### 3.2 Subjective Evaluation

Next, we sought to assess whether pathologists may use a CPP as a tool in BMA cytology. Hematopathologists were asked to decide whether a shown synopsis belongs to the shown CPP or not. Among all 60 slides, the real synopsis report was shown in 26 cases (43.33%). In 46 out of 60 cases (76.67%), pathologists correctly decided whether the CPP matches the shown synopsis or not, as shown in the confusion matrix shown in Figure 6. It is important to note that pathologic evaluation is a subjective task, particularly BMA cytology, so one pathologist’s interpretation may differ from another. Additionally, randomly selected synopses were considered to be negative cases. Therefore, some of those randomly selected synopses might have been semantically similar to the real synopsis. Upon examination of the 5 false positive cases (random but deemed match), it was found that in all these cases, the shown synopsis was similar to the real synopsis. Consequently, the accuracy could be adjusted to 85% with no false positive cases.

**Figure 6:**
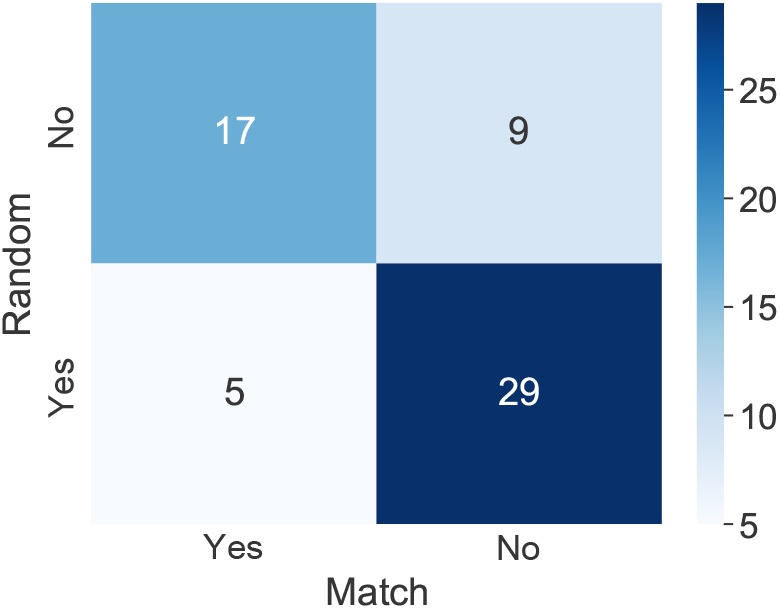
The confusion matrix for the synopsis matching experience. Pathologists were asked to decide whether a displayed synopsis is for the shown CPP or not.

## 4 Discussion

Here, we present a new visualization scheme (CPP) to efficiently and compactly represent the information gathered by an objection detection model from BMA WSI. The usability of CPPs were evaluated by hematopathologists through a synopsis-matching experiment. This evaluation took place in a medically relevant setting. However, we did not provide any additional patient information to the pathologist other than the CPP itself (as a summary of the WSI). This limited amount of information makes diagnosis particularly difficult. In real-world clinical practice, pathology, especially hematopathology, is multimodel, where significant ancillary data, such as clinical history, flow cytometry, and molecular data are required to make a final diagnosis [23]. However, our evaluating pathologists did not have access to data beyond morphology in this type of controlled evaluation. Additionally, the task of BMA reporting is subjective in nature. Thus, two pathologists could have different interpretations of the same BMA slide. Therefore, even if both the digitization process and the AI model were perfect, it would not be expected that the evaluating pathologist to completely agree on a slide synopsis, which was deemed as ground truth in this study. However, at the same time, they would not be expected to differ significantly. Additionally, in our study, there were 5 randomly selected synopses that were falsely deemed matching to the CPP. However, upon examining these shown synopses with the actual ones, we found that the shown and actual synopses were quite similar, with a similar primary diagnosis. Therefore, it can be said that they are true positive cases rather than false positive cases, which could increase the synopsis matching accuracy to 85%.

AI’s lack of explainability is an obstacle to its acceptance for clinical use [24, 25, 26]. It is argued that the current post-hoc explainability approaches might be useful at patient-level decision-making [27]. Therefore, explainability could be addressed while designing the AI workflow implementation. Using CPPs, pathologists were able to assess the quality and clinical relevance of information presented by an AI model. Therefore, such implementation of AI is more explainable as opposed to an AI model making a diagnostic prediction per se. Additionally, one of the common concerns among pathologists is the possibility of their jobs being replaced by AI [28]. CPPs are non-threatening to pathologists as they keep them involved in the model’s decision-making process, potentially acting as a clinical decision-support tool.

In summary, the CPP approach is at the very least a useful tool to efficiently provide pathologists with the information obtained from object detection models in cytology. Additionally, CPPs coupled with reliable object detection models provide automatically generated supplementary information that could potentially augment the current diagnostic workflow. CPPs can be considered a preliminary step towards human-centered AI in cytopathology. CPP may also be used to make cytology reports more understandable for clinical stakeholders other than pathologists. As CPPs are more transferable than WSIs, pathologists could annotate their findings on CPPs and share them with other medical professionals, as shown in Figure 7. Further investigation is required to measure the impact of CPPs if being used as a tool alongside WSIs to augment the digital cytology workflow.

**Figure 7:**
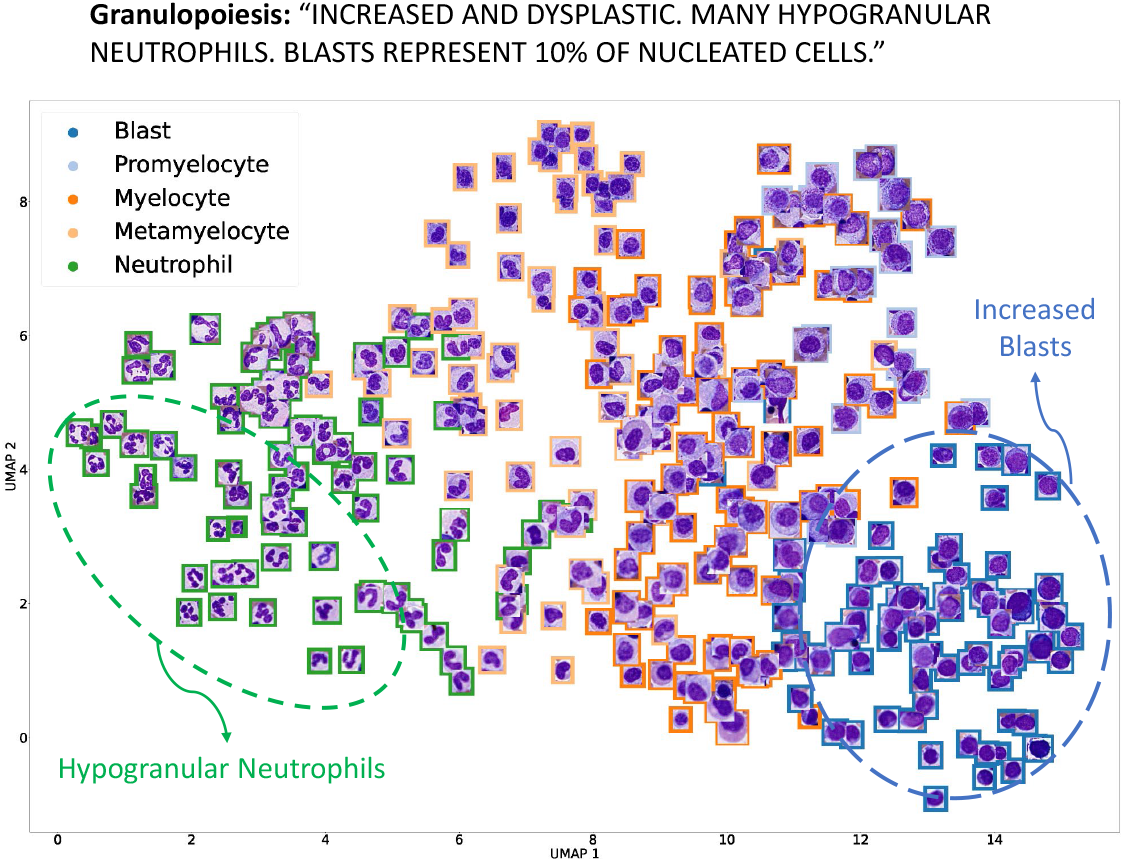
Pathologist could annotate their findings on CPPs which are more transferable that multigigabyte-sized WSIs.

## 5 Data Availability

The WSIs that support the findings of this study are not publicly available due to them containing information that could compromise research participant privacy/consent.

## 6 Code Availability

Code from this study is available at https://github.com/C-Campbell-Lab.

## 7 Competing Interests

The authors declare they are founders of the medical image software startup Parapixel Diagnostics.

## 8 Funding

Funding for this work was provided by a Canadian Cancer Society BC Sparks Grant.

## 9 Author contributions

TD designed and conducted experiments analyzed data, and wrote the paper; HRT designed experiments, analyzed data, provided conceptual input, and contributed to writing the paper; YM provided conducted experiments, designed the software, and contributed to writing the paper; CJVC designed experiments, analyzed data, provided conceptual input and contributed to writing the paper. CJVC, CR, and MS reviewed and evaluated CPPs.

